# An integrated analysis of myeloid cells identifies gaps in *in vitro* models of *in vivo* biology

**DOI:** 10.1101/719237

**Authors:** Nadia Rajab, Paul W Angel, Yidi Deng, Jennifer Gu, Vanta Jameson, Mariola Kurowska-Stolarska, Simon Milling, Chris M Pacheco, Matt Rutar, Andrew L Laslett, Kim Anh Lê Cao, Jarny Choi, Christine A Wells

## Abstract

The Stemformatics myeloid atlas is an integrated transcriptome atlas of human macrophages and dendritic cells that systematically compares freshly isolated tissue-resident, cultured, and stem-cell derived myeloid cell types. We identified two broad classes of tissue-resident macrophages with lung, gut and tumour-associated macrophages most similar to monocytes. Microglia, Kupffer cells and synovial macrophages shared similar profiles with each other, and with cultured macrophages. Pluripotent stem cell-derived macrophages were not reminiscent of fetal-derived cells. Instead, they were characterized by atypical expression of collagen and a highly efferocytotic phenotype. Likewise, Flt3L-derived cord blood dendritic cells were distinct from conventional dendritic cell subsets isolated from primary tissues and lacked expression of key pattern recognition receptors. Myeloid subsets were reproducible across different experimental series, showing the resource is a robust reference for new data. External users can annotate and benchmark their own samples, including annotation of myeloid single cell data at www.stemformatics.org/atlas/myeloid/.

## Introduction

Macrophages are innate immune cells that are found resident in every tissue, with roles in tissue homeostasis, and response to infection or injury. The distinct functional roles of tissue macrophages are reflected in their transcriptional phenotypes: atlases of tissue-resident mouse macrophages, for example, have given great insight into their complexity and heterogeneity (Gautier et al., 2012; Gosselin et al., 2017). Individual transcriptome studies have played an essential role in unravelling the importance of the environment on macrophage phenotype and function (reviewed by (Huang and Wells, 2014)). These data evidence the shared molecular pathways of myeloid cells responding to pathogenic challenge, and the receptiveness of macrophages to environmental cues.

Much of our understanding of macrophage biology, including many of the molecular mechanisms of innate immune signaling, have arisen from mouse gene knock-out studies. However, cross-species comparisons of immune cells highlight differences between mouse and man. These include the glycolytic switch associated with metabolic reprogramming in activated mouse macrophages (Vijayan et al., 2019); divergent patterns of pathogen receptor expression (Vijayan et al., 2012)and transcriptional responses to innate immune stimuli (Schroder et al., 2012). Cross-species comparisons are further hampered by the absence of population level immune-activation maps, with most mouse studies in macrophage biology conducted on a limited number of inbred lines.

The need for improved molecular models of primary human cells is evident from the rising popularity of single cell transcriptomic atlases, exemplified by the human cell atlas consortium (Hay et al., 2018; Regev et al., 2017). However, unbiased profiling of cells also requires computational predictions of cell identity, raising further questions about how best to accurately identify immune cell populations resident in tissues, and discriminate these from circulating or infiltrating peripheral blood cells. The isolation, and identification of tissue-resident myeloid cells can be particularly fraught if populations are rare or hard to isolate using enzymatic or other dissociation methods. These procedures can alter myeloid transcriptomes (Gosselin et al., 2017), resulting in underrepresentation or phenotypic ambiguity of resident macrophages in single cell maps of a tissue. It might be argued that human macrophages suffer from an identity crisis, relying on equivalency to laboratory models of human macrophage biology such as *ex-vivo* culture of monocyte-derived macrophages, which may not be appropriate as a benchmark for specialized tissue functions.

The potential to model tissue residency, disease phenotypes and activation status of human macrophages using pluripotent stem cells (PSC) is a growing area of interest (Reviewed (Lee et al., 2018; Rajab et al., 2018)). However, the anatomical context or developmental ontogeny of these cells is still not well understood, nor their capacity to model specialized behaviors of myeloid cells including roles within a tissue niche. While databases such as BloodSpot (Bagger et al., 2016) and Haematlas (Watkins et al., 2009) provide a useful snapshot of gene expression of blood types, these lack depth with regard to tissue representation, common laboratory models or activating stimuli. Consequently, comparisons of new models of human macrophage biology rely on ad hoc comparisons that do not adequately benchmark their similarity to the diversity of possible macrophage phenotypes. Our motivation, therefore, was to construct a reference atlas of human myeloid biology that draws on important studies already in the public domain, and that can be added to by the research community as new cell models or new profiling platforms become available. Here, we describe an integrated myeloid transcriptome atlas to identify, benchmark and analyze human myeloid subpopulations from *ex vivo, in vivo* and *in vitro* sources. The atlas is made available as an interactive online resource https://www.stemformatics.org/atlas/myeloid.

### A reference atlas for human myeloid biology

We first compiled a reference transcriptional atlas (Figure 1A and Table S1) from public and proprietary transcriptomic data from 44 studies and ~900 samples representing peripheral blood monocytes, tissue-resident, *ex vivo* and *in vitro*-derived macrophages and dendritic cells. Samples were curated with respect to phenotype, source and isolation method. Datasets were processed through the Stemformatics pipeline which includes a stringent set of quality control requirements for hosting on the Stemformatics.org portal (Choi et al., 2019a). We constructed the atlas by implementing a two-step process (Angel et al., 2020). Firstly, transformation of expression values from the original studies to percentile values to facilitate the comparison of different experimental series was performed. Secondly, using a univariate estimation of their platform dependence, genes whose expression values were significantly impacted by the way that they were measured were removed from the atlas. This approach led to reproducible clustering of distinct myeloid classes on a PCA (Figure 1, Figure S1A). The reproducibility of myeloid subsets including dendritic cells, monocytes and neutrophils was validated by projecting an independent RNAseq dataset of well annotated blood cell types from Haemopedia (Choi et al., 2019b) (Figure 1A). Variables such as progenitor source (hematopoietic progenitor cell-, myeloid-, or pluripotent stem cell (PSC)-derived); gene expression (Figure 1B) or culture status (Figure 1C) contributed to clustering of samples. Patterns of gene expression allow users to assess markers of myeloid subsets, such as the expression of *TREM1* on monocytes, monocyte-derived macrophages (MDM), alveolar macrophages and CD1c+ dendritic cells (DC2) (Figure 1B and Table 1).

**Figure 1:**
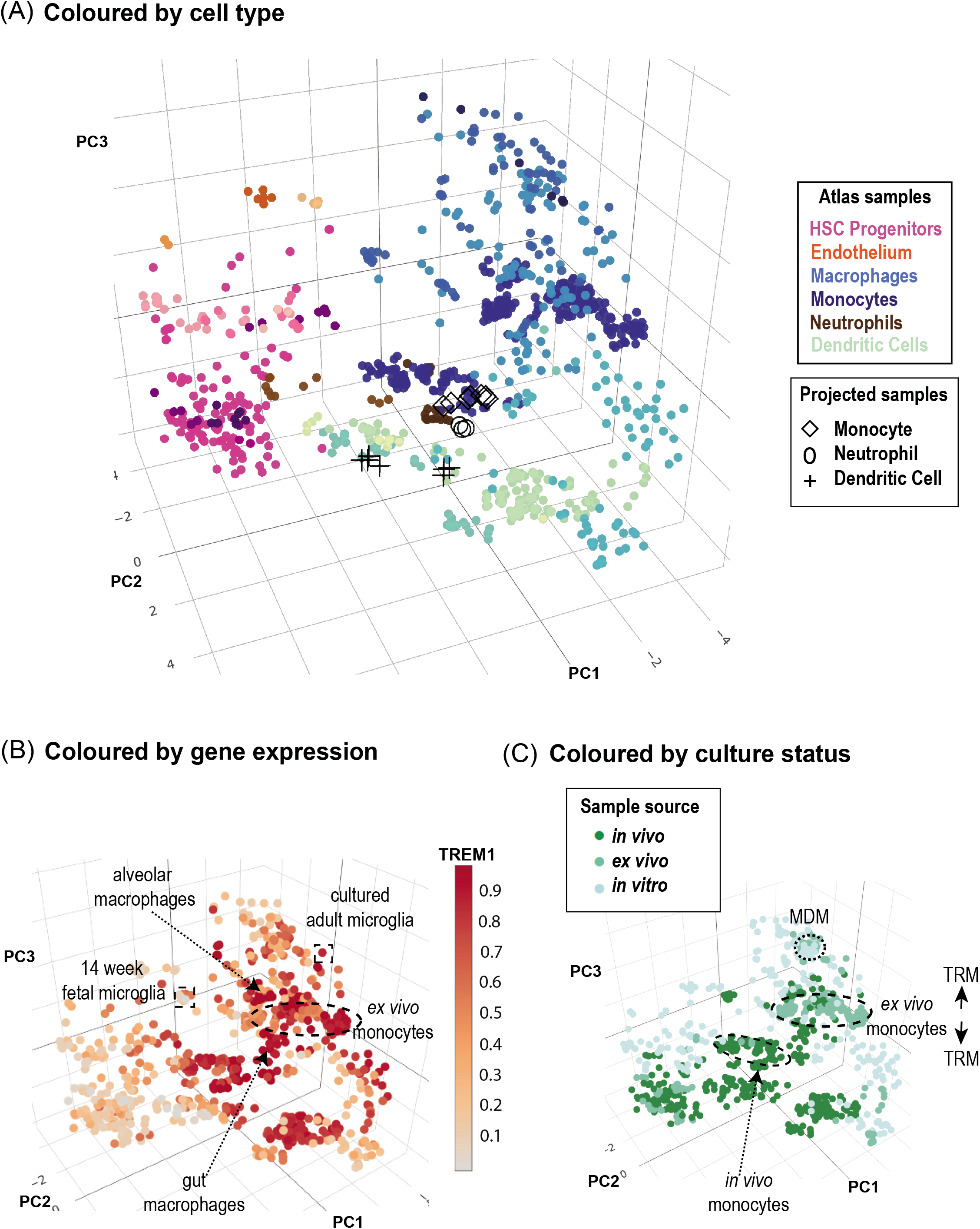
A reference atlas for human myeloid biology. (A) Stemformatics myeloid atlas with samples coloured by cell type. Navy blue - monocytes, blue - macrophages, aqua -Dendritic cells, dark green - CD141+ DC, light green - CD1c+ DC, yellow - pDC, brown – granulocytes, pink stem and progenitor cells, hemogenic endothelium. Validation with Haemopedia RNAseq myeloid samples: diamond shape – monocytes, circle granulocytes, cross DC. (B) Atlas coloured by ranked expression of TMEM1. (Scale bar: high ranked expression (dark red) to low ranked expression (grey) indicating axis of activation) (C) Atlas coloured by culture status (sample source). See also Table 1, Figure S1, and Table S1.

**Table 1:** Atlas inclusion of tissue-resident monocytes and macrophages

| Tissue | Tier 1 (sample #) | Tier 2 | Cell Type |
| --- | --- | --- | --- |
| Blood | *in vivo* (64)<br>*ex vivo* (162) | myeloid | monocyte |
| Cord Blood | *ex vivo* (51) | myeloid | monocyte |
| Gut | *in vivo* (15) | myeloid | macrophage |
| Synovium | *in vivo* (18) | myeloid | macrophage |
| Brain | *in vivo* (10)<br>*ex vivo* (21) | myeloid | microglia |
| Lung | *in vivo* (14) | myeloid | macrophage |
| Liver | *ex vivo* (5) | myeloid | Kupffer cell |
| Ovarian Tumour | *in vivo* (4) | myeloid | macrophage |

### Two broad classes of tissue resident macrophage span either side of a monocyte border

Isolation of primary tissue-resident macrophages is particularly difficult as this can result in alterations in phenotype (Gosselin et al., 2017). The difficulty of isolating tissue-resident populations from healthy human tissue is evident from the spread of tissue resident macrophages in comparison to tissue-resident dendritic cells (Figure 1), noting that several of the macrophage datasets were obtained through surgical biopsies from patients with inflammatory disease, and indeed mapped across the inflammatory axis. Tissue resident macrophages, including Kupffer cells, microglia, alveolar macrophages, gut and synovial macrophages occupy a broad niche on the atlas between dendritic cells, peripheral blood monocytes, and cultured monocytes (Figure 1 and Figure S1). Although there was no discrete partitioning of macrophages from individual tissues, distinct classes of tissue resident macrophages were observed in the atlas – (the alveolar, colon and macrophages isolated from tumour ascites (TAM) (Figure S1C) grouped together between cultured monocytes and CD1c+ dendritic cells, and a second spread containing synovial macrophages, microglia and Kupffer cells aligned with the iPSC-derived cells.

### DC differentiated from cord blood lack an *in vivo* equivalent

Circulating and tissue resident dendritic cells (DCs) occupy a distinct transcriptional niche from monocytes or macrophages. *In vitro*-differentiated dendritic cells, expanded from cord blood progenitors with FLT3L, do not resemble *in vivo* conventional DCs (DC1 or DC2; Figure 1B, Figure 2A), but are part of an activation axis shared by DC and macrophages. The hallmark of *in vitro*-DCs are low expression of receptors such as *CX3CR1, IL18R1* and *TLR7* (Figure 2B, 2C and Table S2). Other molecules, such as the cell-fusion protein *DC-STAMP* are gained in culture (Figure 2C and Table S2). These cells are also closely associated with MDMs, which may contribute to some of the confusion in the literature about their ontogeny. It is unlikely that this is due to the cord blood origins of the majority of these datasets, as CD45+ cells isolated from cord blood engrafted into humanized mice are also included in the atlas (Minoda et al., 2017) and these recapitulate *in vivo* DC1 and DC2 transcriptional phenotypes.

**Figure 2:**
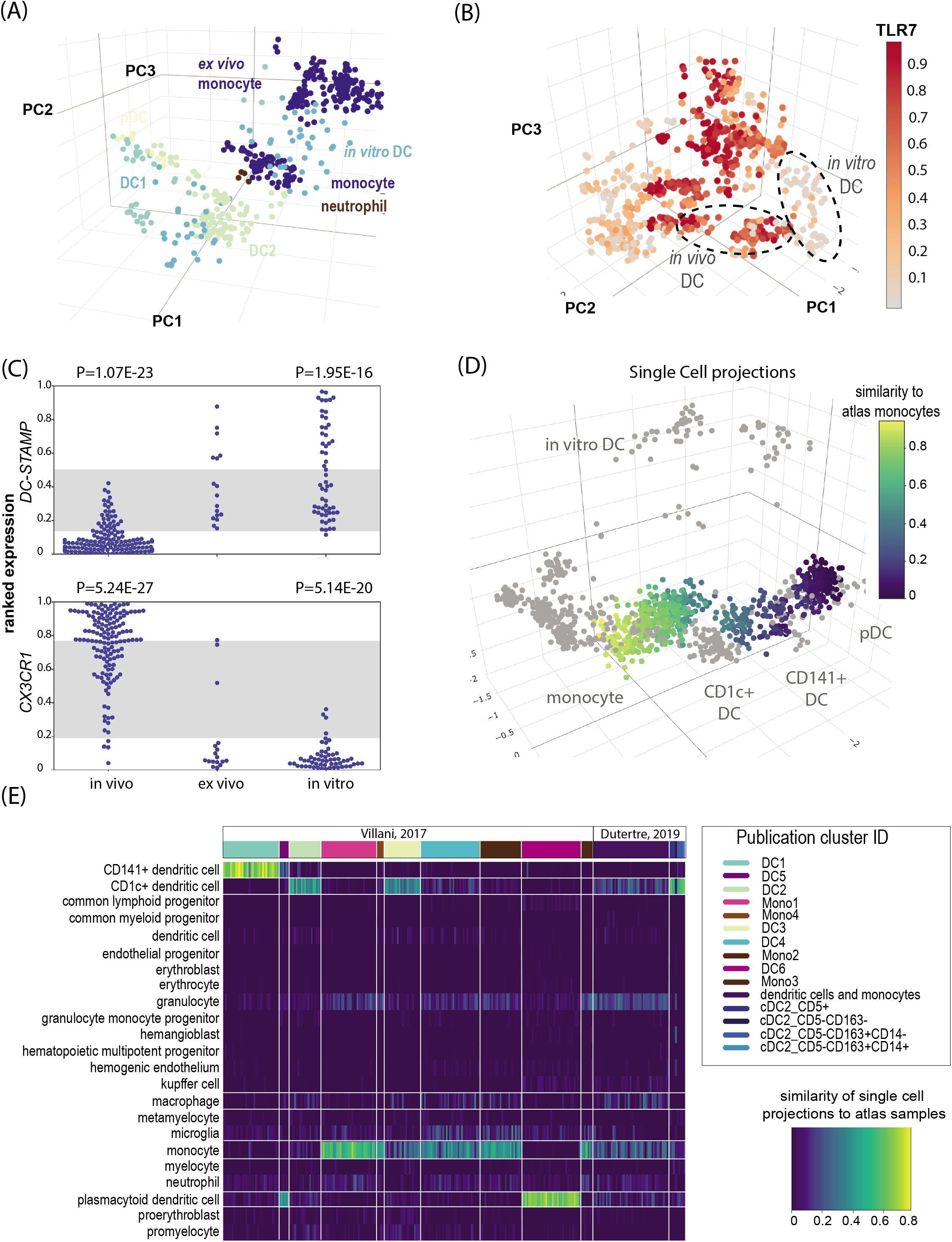
Cultured and *in vitro*-derived dendritic cells do not capture aspects *in vivo* myeloid biology. (A) DC subsets displayed in the atlas – aqua *in vitro* (cord blood derived) DC, dark green DC1 (CD141+), light green DC2 (CD1c+), yellow - pDC, brown neutrophils (B) atlas coloured by ranked expression of TLR7. (Scale bar: high ranked expression (dark red) to low ranked expression (grey)). (C) Ranked expression (Y-axis) of receptor CX3CR1 and cell-fusion protein DC-STAMP *in vivo* dendritic cells (n=145), *ex vivo* dendritic cells (n=17) and *in vitro*-derived dendritic cells (n=57). Grey stripe indicates variance attributable to platform. P-value: Mann-Whitney-Wilcoxon rank sum test. (D) Single cell projections of Villani *et al*. (2017) and Dutertre *et al*. (2019) samples onto atlas (F) Heatmap derived from Capybara analysis of Villani *et al*. (2017) and Dutertre *et al*. (2019) samples compared to myeloid cell types. Colour gradients reflect similarity of single cell clusters to atlas cell types (dark least similar, to light most similar). See also Figure S2 and Table S2.

The cDCs form distinct clusters along the activation axis of the atlas. This includes blood samples taken from donors after vaccination (Banchereau et al., 2014) or cells isolated from inflammatory fluids (Segura et al., 2013) and we speculate that these represent distinct and reproducible activation phenotypes. To further evaluate DC subtypes we projected two single cell RNAseq datasets describing blood monocytes and dendritic cells (Villani et al., 2017) and (Dutertre et al., 2019) using the myeloid atlas as the reference (Figure 2D, Figure S2). We confirmed the original observation of a new *AXL*+ *SIGLEC6*+ (AS-DC) subset that shared classical cell surface markers with plasmacytoid DCs (pDCs) and DC1s (Figure 2E). AS-DCs were suggested to be the subset contaminating traditional pDC isolation strategies, responsible for observations that pDC can stimulate T-cells (Villani et al., 2017). There has been some dispute between the Villani and Dutertre studies on the identity of DC2 subsets (Dutertre et al., 2019). An advantage of projecting both datasets to the same reference is that the subtypes can be evaluated against a broader set of reference cell types. Here, the Dutertre DC2 subsets aligned closely with atlas DC2 cells, while the spread of Villani clusters was much greater, and clearly aligning to respective DC and monocyte groups on the atlas. Annotation with a Capybara similarity estimate (Figure 2E) predicted that in addition to the 2 distinct plasmacytoid DC (pDC) subsets in the Villani data, three distinct DC2 (CD1c+) subsets were present, including an intermediate DC2 subset that sat between the classical monocyte and DC2 areas (Figure 2E, 2F and Figure S2). Given the number of additional DC1 and DC2 subsets observed on the myeloid atlas, we predict that further functional characterization of activated DC phenotypes is warranted.

### Monocytes rapidly adapt to culture

Monocytes are post-mitotic blood cells derived from bone marrow progenitors that are short lived in circulation and can repopulate macrophages in some tissue niches. The largest population of circulating monocytes is marked by high levels of expression of the LPS co-receptor CD14, which is typically used to isolate monocytes from blood. Intermediate and nonclassical subsets are marked by acquisition of the type III FcRƔ, CD16 (Schmidl et al., 2014) and are included in the atlas. Cultured monocytes have been previously described as ‘activated’, but while we observe a distinct culture phenotype (Figure 3), the transcriptome of cultured cells mimics many of the features of a monocyte after extravasation into tissue. Figure 3B shows the grouping of peripheral blood monocytes, in a distinct cluster to monocytes that have been exposed to tissue culture plastic and culture media. This culture phenotype is typified here by a decrease in endothelial-adhesion proteins including the selectin *SELL* (Figure 3D). Regulators of RAS/RAF signaling including *SPRED2* (Wakioka et al., 2001) have an elevated expression in cultured monocytes (Figure 3E), consistent with spreading and migration across tissue culture plastics. The classical MDM requires *in vitro* differentiation of monocytes using several days exposure to growth factors such as macrophage-colony stimulating factor (M-CSF; CSF-1) or granulocyte-macrophage colony-stimulating factor (GM-CSF). These group distinctly from the cultured monocyte cluster, spreading further upwards along the culture axis (Figure 3F).

**Figure 3:**
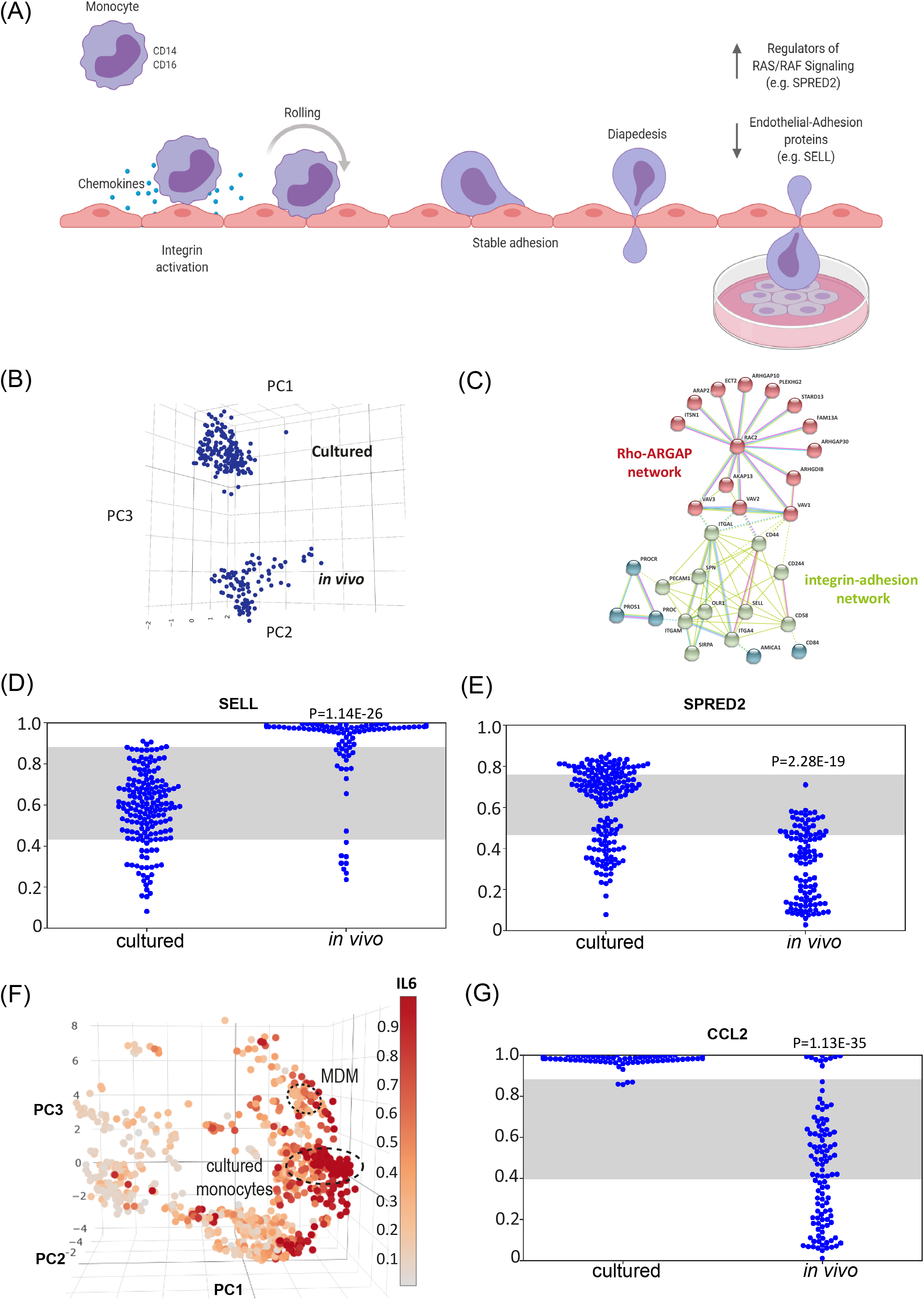
Monocytes acquire a culture phenotype. (A) Schematic of rolling monocytes, highlighting cultured cells mimic many of the features of a monocyte after extravasation (B) Cultured monocytes form a distinct cluster away from *in vivo* monocytes along PC3 (C) STRING-DB network of top-ranked genes differentially expressed between peripheral blood (*in vivo*, n=107) and cultured monocytes (*ex vivo*, n=171) indicating upregulation of cytoskeletal proteins and down regulation of endothelial-adhesion proteins (D) Ranked expression (Y-axis) of gene involved in endothelial adhesion, SELL, comparing cultured monocytes (n=171) with monocytes directly profiled from blood (*in vivo*, n=107). Grey stripe indicates variance attributable to platform. P-value: Mann-Whitney-Wilcoxon rank sum test (E)Ranked expression (Y-axis) of gene involved in the regulation of RAS/RAF signaling comparing cultured monocytes (n=171) with monocytes directly profiled from blood (*in vivo*, n=107). Grey stripe indicates variance attributable to platform. P-value: Mann-Whitney-Wilcoxon rank sum test (F) atlas coloured by ranked expression of IL-6 (Scale bar: high ranked expression (dark red) to low ranked expression (grey) indicating axis of activation) (G) Ranked expression (Y-axis) of Chemokine CCL2 comparing cultured monocytes (n=171) with monocytes directly profiled from blood (*in vivo*, n=107). Grey stripe indicates variance attributable to platform. P-value: Mann-Whitney-Wilcoxon rank sum test. See also Table S3.

The culture phenotype acquired by monocytes appears to be a prelude to activation, which can be observed along an adjacent axis in Figure 3F and is exemplified by the expression of *IL6*. Pathogen-activated phenotypes of cultured monocytes are typified by high expression of this and other cytokines. We observe that cultured monocytes express higher levels of chemokines, such as *CCL2*, than peripheral monocytes (Figure 3G). Furthermore, we find that culture induces the expression of *SLAMF1* (Table S3), which has shown to be necessary for TLR4 activation in human macrophages (Yurchenko et al., 2018). Moreover, this culture-phenotype is further exemplified by higher levels of *ITGB8* than circulating monocytes (Table S3), which is necessary for activating latent TGF-β (Kelly et al., 2018)

### Pluripotent stem cell-derived macrophages do not recapitulate hematopoietic ontogenies

Many PSC-derived systems recapitulate fetal, rather than adult phenotypes, so it is no surprise that others have argued that PSC-derivation protocols mimic primitive rather than definitive myeloid biology. This is largely based on *MYB* expression, which is associated with definitive hematopoiesis and has high expression in hematopoietic progenitor cells isolated from bone marrow. It is clear that *MYB* is not required for PSC-derived myelopoiesis as macrophages can be grown from *MYB* knock-out embryonic stem cells (Buchrieser et al., 2017). Nevertheless, *MYB* is highly and ubiquitously expressed in PSC-myeloid progenitors, including common myeloid progenitors and hemogenic endothelium differentiated from pluripotent stem cell, and indeed, *MYB* is retained at high levels in some PSC-derived microglia (Figure 4A).

**Figure 4:**
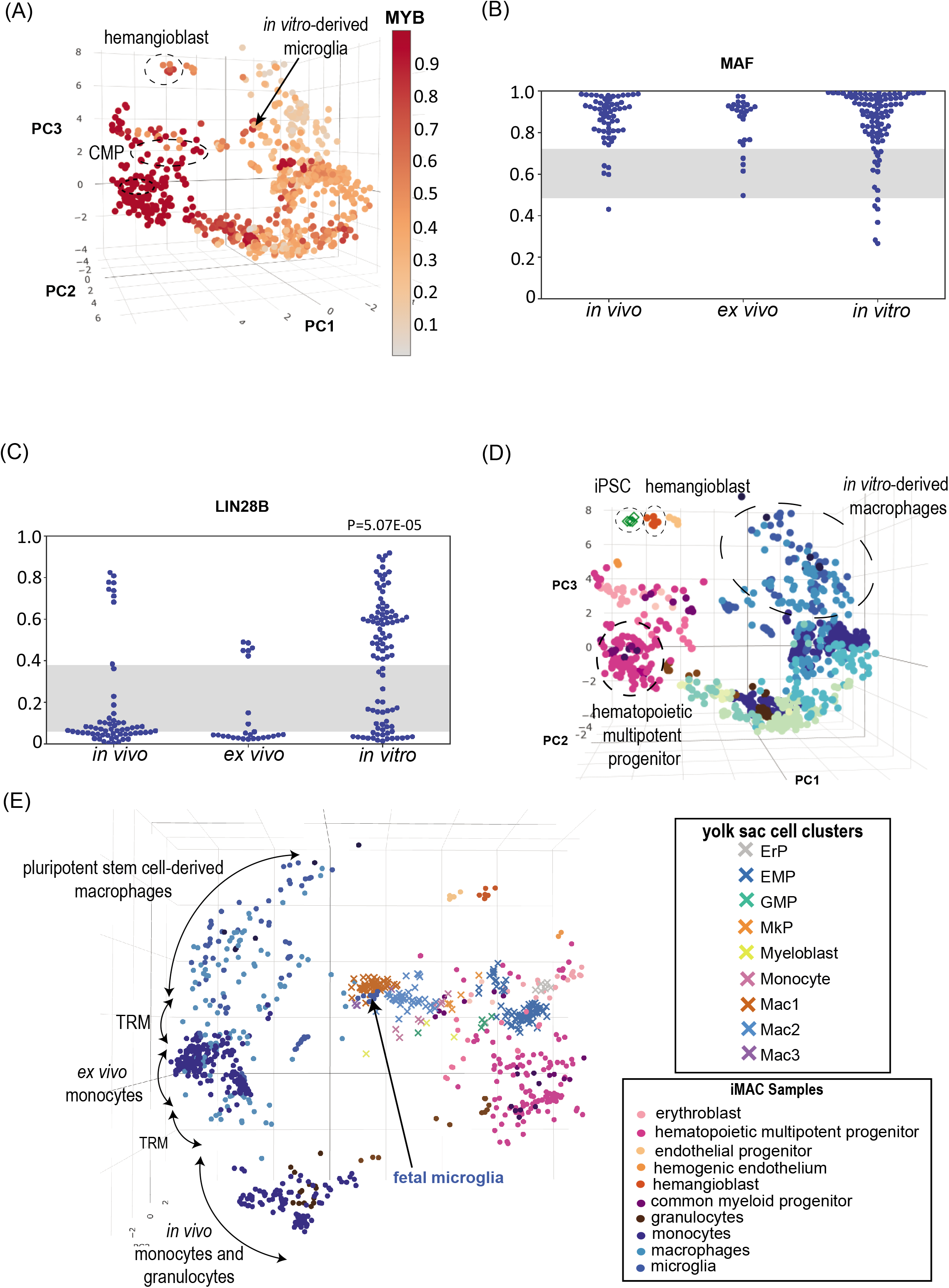
Pluripotent stem cell-derived macrophages do not recapitulate hematopoietic ontogenies. (A) atlas coloured by ranked expression of MYB (Scale bar: high ranked expression (dark red) to low ranked expression (grey). (B) Ranked expression (Y-axis) of MAF expression from *in vivo* (n=61), *ex vivo* (n=26) and *in vitro*- (n=96) derived macrophages (gut, synovial, kupffer, microglia, macrophage). Grey stripe indicates variance attributable to platform. (C) Ranked expression (Y-axis) of LIN28B expression from *in vivo* (n=61), *ex vivo* (n=26) and *in vitro*- (n=96) derived macrophages (gut, synovial, kupffer, microglia, macrophage). Grey stripe indicates variance attributable to platform. P-value MannWhitney-Wilcoxon rank sum test. (D) atlas with samples coloured by cell type with projection of iPSC samples highlighting their position in comparison to *in vitro*-derived macrophages, hematopoietic multipotent progenitors and hemangioblast. (E) Single cell projection of (Bian et al., 2020) human fetal yolk sac cell clusters onto the atlas following an 8 cell aggregation. See also Figure S3 and Table S4 and Table S5.

PSC-macrophage studies that were overlaid onto the atlas grouped broadly with cultured monocytes and tissue-resident macrophages. Despite arguments on recapitulation of fetal origin, PSC-macrophages, including PSC-microglia and PSC-Kupffer cells, form an extended group associated with high expression of the human homologue of the *F4/80* antigen, *ADGRE1*, as well as high expression of lipid-scavenging receptors such as *SCARB1* (Table S4). MAF expression is indistinguishable in macrophages of different origin (Figure 4B). However, *LIN28B* is highly expressed in *in vitro*-derived macrophages compared to *in vivo* and *ex vivo* cells (Figure 4C), which may point to incomplete silencing of the Let7 microRNA pathway and maintenance of a fetal state (Grant Rowe et al., 2016; Zhou et al., 2017), but which is also characteristic of myeloid leukemias.

When pluripotent stem cells are projected onto the atlas, we can see a differentiation trajectory that is orthogonal to the hematopoietic multipotent progenitor trajectory (Figure 4D). The differentiation trajectory from pluripotent stem cell to macrophage aligns more closely with an endothelial progenitor than the hematopoietic intermediates observed along PC1 of the atlas. Projection of primitive hematopoietic progenitors from Carnegie staged embryos at CS11 to CS23 show a high degree of similarity between definitive and primitive differentiation trajectories, with a predominant erythroid overlap in early yolk sac progenitors, followed by rapid emergence of myeloid progenitors, and primitive macrophages. These do not overlap with the PSC-axis, but do clearly sit with similar late fetal microglia (20 week old, (Thion et al., 2018)) and adult bone-marrow equivalents (Watkins et al., 2009) (Figure 4E and Figure S3).

Some other phenotypes previously attributed to ontogeny in PSC-derived cells may rather reflect a more general culture context. For example, *ADGRE1 (F4/80)* expression has been attributed to yolk-sac derived myeloid cells in mouse (Schulz et al., 2012). While high on PSC-derived cells, *ADGRE1* is also clearly upregulated in *ex vivo* cultured macrophages. This is exemplified by primary human microglia, which have low expression of *ADGRE1* in comparison to *ex vivo* culture or PSC-derived cells (Table S4 and Table S5).

### Pluripotent stem cell-derived macrophages share features with tissue-resident macrophages despite poor maturation

Macrophages derived from human PSCs offer new opportunities to model *in vivo* macrophage biology. When reviewing the studies contributing to this atlas, we noted that PSC-derived macrophages are typically benchmarked against monocyte-derived macrophages (MDMs), or cultured primary cells, using a suite of phenotyping techniques. Each experiment includes a small number of samples for transcriptional profiling, with a few notable exceptions (Alasoo et al., 2018). We argue that, given the spectrum of possible resident tissue macrophage phenotypes, it would be more useful to compare PSC-derived cells against an atlas of possible macrophage phenotypes. Whilst several groups reuse publicly available tissue macrophage data, the opportunity to carry out large-scale comparisons to different primary myeloid cells has been limited by the availability of relevant data on a compatible platform.

Primary microglia included in the atlas include both *in vivo* isolated fetal and cultured *ex vivo* fetal and adult microglia. The profiles of *in vivo* isolated fetal microglia cluster apart from the spread of *ex vivo* cultured adult and fetal microglia (Figure S1B). Projection of single cell data from human fetal yolk sac and head (Bian et al., 2020) show high concordance with the atlas fetal microglia data, suggesting that age and culture environment override an ontogeny signature (Figure 4 and Figure S3). Cultured tissue macrophages including *ex vivo* primary microglia or Kupffer cells shared a broad transcriptional signature with MDMs, and pluripotent-stem cell derived myeloid cells. Microglia represent just over a third of PSC-directed myeloid differentiation studies in the atlas. These do not resolve into a unique cluster but share transcriptional phenotypes with PSC-derived and tissue-resident macrophages

There are exceptions which include some ‘cytokine-matured’ PSC-derived microglia samples from (Abud et al., 2017). These are close to the *in vivo* fetal microglia samples of the PSC-microglia but are also closely associated with other primary tissue resident macrophages from lung, joint and gut. The atlas does provide an opportunity to review the expression of markers thought to distinguish microglia from other primary macrophages. *TMEM119*, for example, is largely restricted to primary or PSC-derived microglia, although some PSC-microglia samples have low expression of this marker (Figure S1B). *P2RY12* is variably expressed across all microglial samples, but its expression is also evident in different tissue-resident samples including those derived from gut and synovial tissues (Figure S1C and Table S4 and Table S5).

The majority of PSC-derived macrophages have low expression of HLA relevant genes including CIITA, a known master regulator of MHCII gene expression, which suggests poor maturation and limited capacity to present antigen to lymphocytes. Nevertheless, some *in vitro*-derived macrophages cultured with stimulating factors (Figure S1D) do show inducible *CIITA* expression, demonstrating that they have the capacity to express antigen-presenting machinery. It is also worth noting additional culture conditions that result in high *CIITA* expression without interferon-stimulation (Figure S1D). This may be the result of long-term culture conditions for microglia samples (Abud et al., 2017), or reflect prior conditioning of myeloid progenitors (Honda-Ozaki et al., 2018).

### Pluripotent stem cell-derived macrophages display transcriptional hallmarks of efferocytosis

Efferocytosis, or apoptotic cell clearance, has broad immunomodulatory effects (Reviewed by (Elliott et al., 2017)) and active engulfment and clearance of cells by PSC-macrophages is clearly observed in the absence of any inflammatory activation (Supplementary Video). Efferocytosis (Figure 5A) modulates macrophages from a pro-inflammatory phenotype to one with resolving qualities (Yamaguchi et al., 2014), consistent with the patterns of gene expression observed in cultured macrophages. A study (Cao et al., 2019) demonstrated higher lipid uptake in PSC-macrophages compared to peripheral blood MDMs, concordant with higher expression of efferocytosis-related genes including *S1PR1* and *MERTK*. We confirm that *MERTK* is generally highly expressed in PSC-derived macrophages, but that there is also a tissue-resident distribution of *MERTK* expression, with very low levels observed in primary alveolar macrophages, and highest levels observed in human fetal microglia (Figure 5B). Tissue-resident macrophages are known first-responders to tissue damage and are key in orchestrating inflammation and its subsequent resolution. This appears to be a phenotype that is selected for in cultured macrophages and may be an inevitable consequence of derivation that strives for high cell yield.

**Figure 5:**
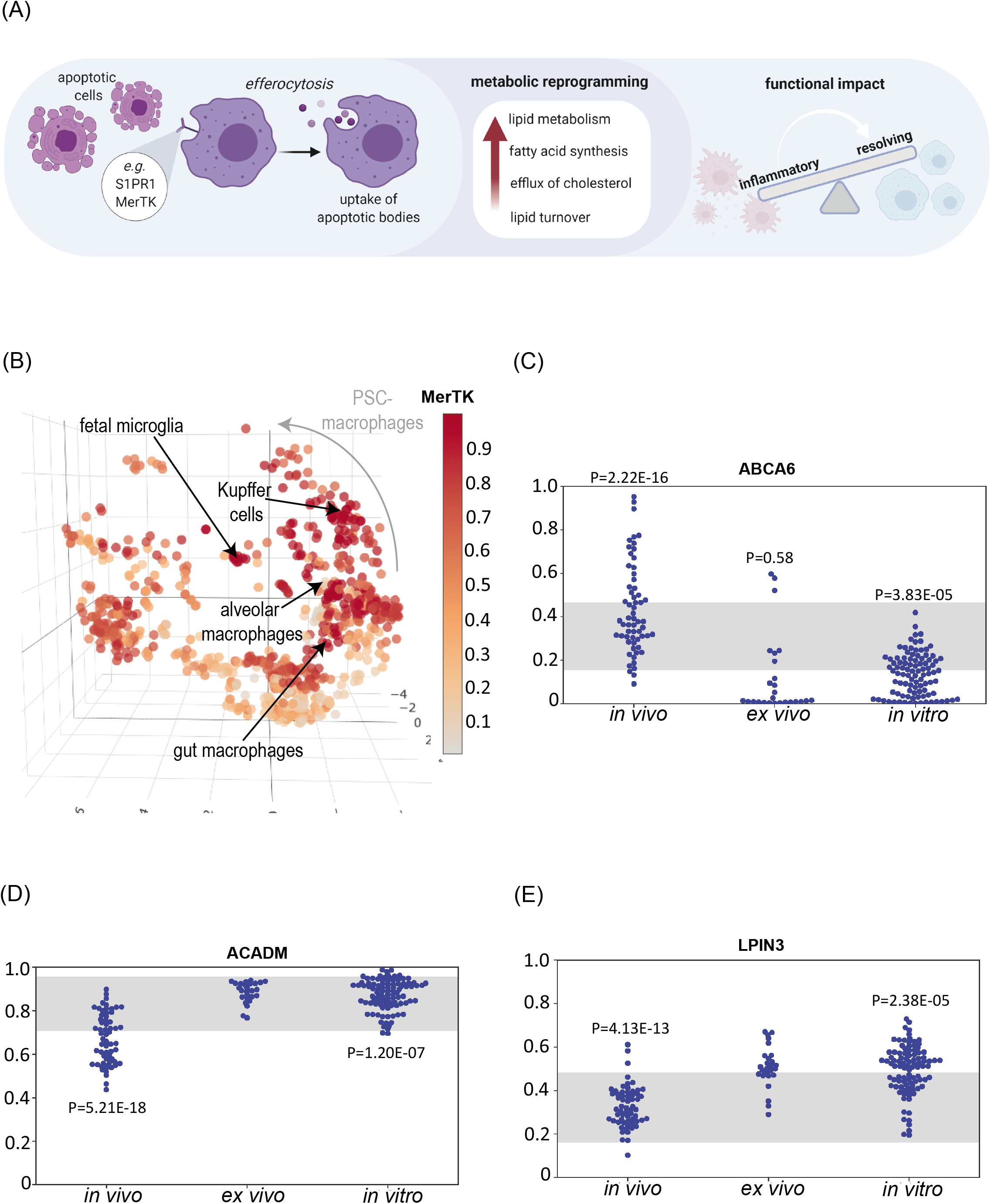
Pluripotent stem cell-derived macrophages display transcriptional hallmarks of efferocytosis. (A)Schematic of impact of efferocytosis on cell metabolic reprogramming and function (B) atlas coloured by ranked expression of MERTK (scale bar: high ranked expression (dark red) to low ranked expression (grey). (C,D,E) Ranked expression (Y-axis) of genes comparing *in vivo* (n=61), *ex vivo* (n=26) and *in vitro*-derived (n=96) macrophages (gut, synovial, kupffer, microglia, macrophage) for (C) cholesterol efflux (D) mitochondrial acyl-CoA dehydrogenase (E) phosphatidate phosphatase. See also Table S4 and Table S5.

Lipid homeostasis is an important role for resident tissue macrophages. A high proportion of genes differentially expressed between *in vitro*-derived macrophages/microglia/kupffer cells and tissue resident cells are involved in lipid transport, catabolism and in buffering the cells from concomitant stresses associated with lipid turn-over. For example, reduced expression of *ABCA6* is consistent with high efflux of cholesterol from these macrophages (Figure 5C). Higher levels of mitochondrial acyl-CoA dehydrogenase *ACADM* (Figure 5D) and phosphatidate phosphatase *LPIN3* (Figure 5E) suggests high lipid turnover.

There has been growing interest in the importance of metabolic reprogramming in macrophage responses, so we asked whether media supplementation could explain the spread of PSC-derived macrophages on the atlas. All PSC-derivation protocols supplement media with fatty- or amino-acids, including L-Glutamine, non-essential amino acids (NEAA), Linoleic and Linolenic acids.

Some methods add fetal bovine or calf serum, but there was no obvious correlation between serum addition and without. Overall, factors are so ubiquitously used that a particular supplement alone could not explain the differences between PSC and cultured primary macrophages.

### Pluripotent stem cell-derived macrophages express high levels of collagen

Collagen production and deposition alongside extracellular matrix remodeling are processes involved in wound healing and scarring. Macrophages are instrumental in instructing tissue repair, particularly through the production of growth factors such as TGF-β, IGF1 and PDGF (Shook et al., 2018). Secreted growth factors drive fibroblasts and endothelial cells to produce extracellular matrix components, promoting keloid formation as well as angiogenesis. This model has macrophages influencing collagen deposition by neighboring stromal cells, however, in some instances are capable of contributing to collagen deposition as demonstrated in mouse and zebrafish injury models (Simões et al., 2020). Gene-set enrichment analysis of the genes that are most correlated with *in vitro*-derived macrophages moving away (Figure 6A) from the tissue resident populations revealed that the most significant pathways in these cells involved collagen synthesis and production (Table S6). A STRING protein-protein interaction network (Figure 6B) shows that this phenotype is significantly enriched for highly connected matrix remodeling, collagen deposition and cadherin-mediated cell-cell and cell-matrix interactions.

**Figure 6:**
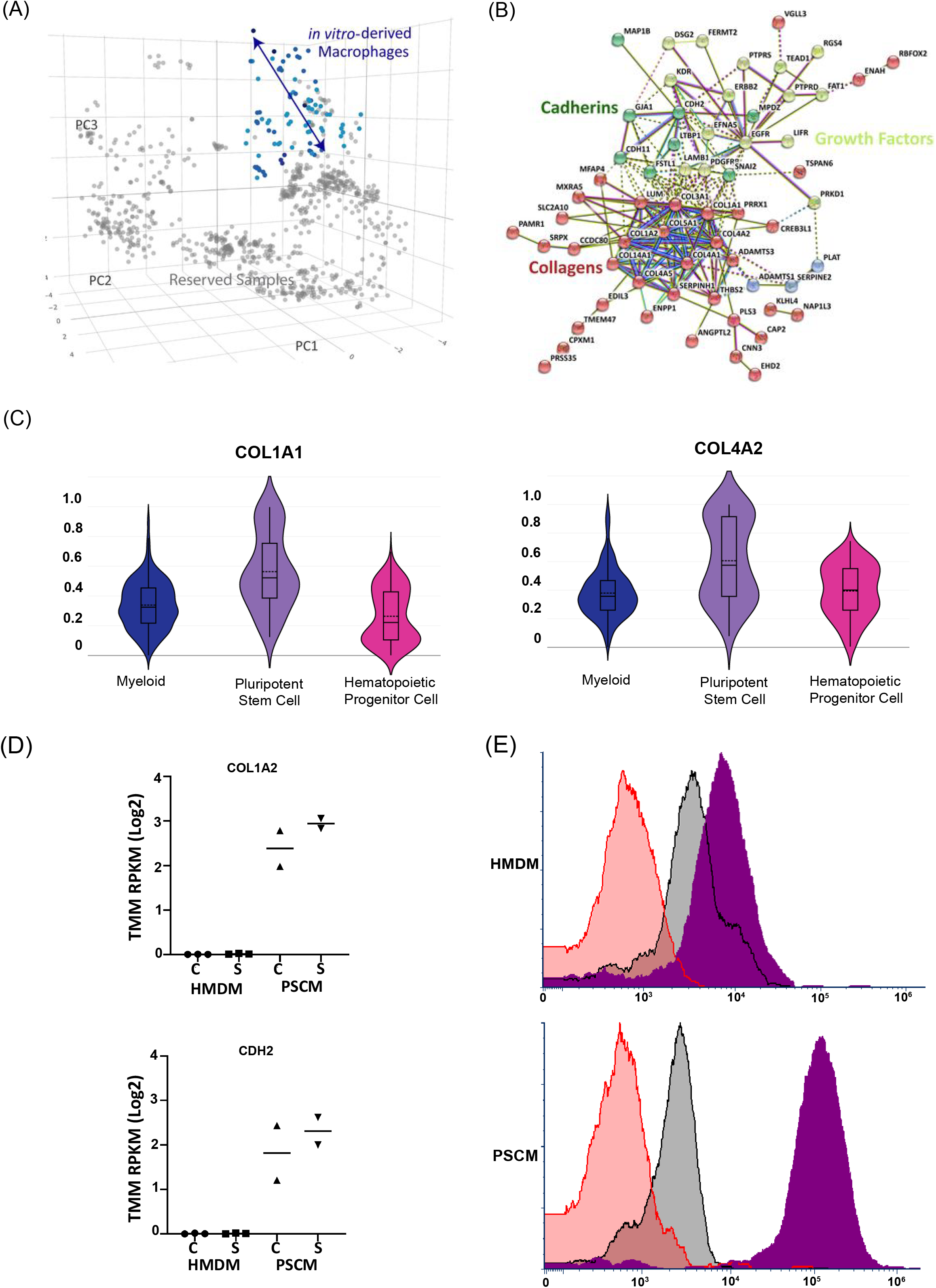
Pluripotent stem cell-derived macrophages express high levels of collagen. (A) atlas coloured by cell source to highlight *in vitro*-derived macrophages used for regression testing (B) STRING_DB Protein-Protein network of *in vitro*-derived macrophages highlights enrichment of collagen, growth factor and cadherin networks. Line color indicates the type of interaction evidence. Light blue solid lines indicate known interactions from curated databases, solid pink line indicates known interactions that have been experimentally determined, bright green lines indicate gene neighborhood predicted interactions, red lines indicate gene fusions predicted interactions, dark blue lines indicate gene co-occurrence predicted interactions, yellow/green lines indicate textmining, black lines indicate co-expression, and light purple lines indicate protein homology. (C) Violin plots of myeloid- (n=584), pluripotent stem cell- (n=116) and hematopoietic progenitor- (n=201) derived cells comparing expression of collagen genes (COL1A1 and COL4A2)(D) mRNA-seq gene expression from human peripheral blood monocyte-derived macrophages (HMDM) (n=3) and human pluripotent stem cell derived macrophages (PSCM) (n=2) samples (C= control, S = stimulated) (E) Intracellular flow cytometry analysis of human peripheral blood monocyte-derived macrophages (HMDM) and human pluripotent stem cell derived macrophages (PSCM), representative of 2 experimental repeats (n=2 HMDM, n=2 PSCM). Red = no primary antibody control, Black = isotype control, Purple = Type I Collagen stain. See also Table S6.

Perhaps pluripotent stem cell-derived cells are being driven to adopt a pro-fibrotic phenotype through the derivation or culture contents. Initial observations on analysis of myeloid-, pluripotent stem cell- and hematopoietic progenitor-derived cells, highlight higher expression of collagen genes in pluripotent stem cell-derived cells (Figure 6C). To investigate whether this phenotype is due to culture impact, we isolated peripheral blood monocytes and derived human PSC-progenitors and cultured these cells with the same culture media in the presence of M-CSF for 5 days to drive macrophage differentiation. On day 5, cells were either stimulated with LPS for 2 hours before extraction or extracted as control samples for sequencing analysis. Analysis of sequencing samples showed the same enriched collagen and cadherin networks with high expression of these genes observed in PSC-macrophages, with little or no expression in monocyte-derived macrophages regardless of whether cells were stimulated or not (Figure 6D). To determine this at a protein level, we carried out intracellular flow cytometry analysis of type I collagen in PSC- and monocyte-derived cells (Figure 6E). Compared with staining controls, we observed higher levels of type I collagen in PSC-macrophages. Collectively, our results suggest that although culture media can impact cell phenotype, it is not the main driver for this pro-fibrotic phenotype in PSC-macrophages in the final stages of differentiation.

## Discussion

Human macrophage biology is integral to the development of homeostasis and disease mechanisms in every tissue in the body, but our understanding of human myeloid biology is limited by the quality by the models available to us. Here we describe an integrated myeloid transcriptome atlas as a novel resource to identify myeloid cells in single cell datasets and to benchmark *in vitro* models of *in vivo* biology. Implementation in Stemformatics enables users to upload their own data to benchmark cell types against the atlas for rapid and intuitive cell-classification. The resource is scalable and will grow as the availability of new tissue resident samples and myeloid models become available.

Transcriptional benchmarking of myeloid subsets, and particularly those benchmarking new derivation methods, typically draw on a small number of reference samples. Standard analysis workflows include normalisation methods that remove technical batch effects, and harmonise the behaviour of samples assigned to the same class. When combining data that confounds technical batch with the biology of interest, this process inevitably overstates similarities within groups and over emphases differences between groups. The result is a somewhat self-fulfilling analysis, with the outcome predicted by the data processing approach. Here, we offer an alternative for benchmarking new models of myeloid biology against a reference atlas constructed from a large number of well phenotyped and curated published data. We demonstrate reproducible classification of major myeloid cell classes, including the influence of culture or derivation method on the phenotype of those cells. Projection of external data into the atlas further demonstrates that these phenotypic patterns are reproducible, even at the resolution of the single cell.

Our analyses highlight that there is room for improvement in the development of *in vitro* model systems that attempt to mimic *in vivo* counterparts. We demonstrate that cord blood-derived dendritic cells differentiated ex vivo from monocytes or CD34 progenitors don’t adequately capture key aspects of *in vivo* myeloid biology. Likewise, by benchmarking curated public data of PSC-macrophages and their precursor cells against the atlas, it is apparent that these represent neither definitive nor primitive myelopoiesis, or rather, that they imperfectly recapitulate aspects of both. PSC-conditions clearly do not mimic the developmental time-frame nor tissue niche of yolk-sac, fetal liver or bone marrow. PSC-macrophages do recapitulate many aspects of ex-vivo cultured tissue macrophages, but there is little evidence for cultured microglia being distinct from any other cultured macrophage. In our hands, PSC-macrophages display transcriptional hallmarks of efferocytosis and surprisingly collagen production, which may suggest that the derivation process of these cells are driving a pro-fibrotic phenotype. These observations offer new opportunities to co-opt additional transcriptional networks that may arise from the tissue niche, to improve stem cell derived myeloid models.

## Supporting information

Video of iPSC-derived macrophages

## Acknowledgements

The authors thank Tyrone Chen and Othmar Korn for assistance with data processing and Isha Nagpal for website development to support the Stemformatics interactive viewer. The authors thank Ramaciotti Centre for Genomics (University of New South Wales; Sydney) and University of Glasgow Polyomics facility for mRNA sequencing, and the Melbourne Cytometry Platform for flow cytometry assistance. This work was funded by Stem Cells Australia, an Australian Research Council Special Research Initiative [SRI110001002] to CAW; NHMRC Synergy [APP1186371] to CAW; Wellcome Trust catalyst funding WELLCOME (097821/Z/11/B) to CAW, SM, MKS; SM work was supported by the Research into Inflammatory Arthritis Centre Versus Arthritis (RACE) [#20298]; MKS is funded by Versus Arthritis UK [#20298 & #22072]. NR is funded by the Centre for Stem Cell Systems and the CSIRO Synthetic Biology Future Science Platform. CAW is funded by a Future Fellowship from the Australian Research Council [FT150100330]. JC is funded by the JEM Research Foundation to the Stem Cell Atlas.

## Author Contributions

Conception NR, JC, CAW; Experimental Investigation and Interpretation NR, VJ, JG, CAW; Experimental Resources ALL, CAW; Methodology PWA, JC, YD; Data provider NR, MKS, SM; Curation NR, MR, CMP, CAW; Statistical analysis YD, KALC, PWA; Writing – original draft NR, CAW; Writing - review and editing NR, PWA, SM, ALL, KALC, JC, CAW; Supervision CAW, ALL, KAC; Project Funding - CAW.

## Supplemental Figures

**Figure S1:**
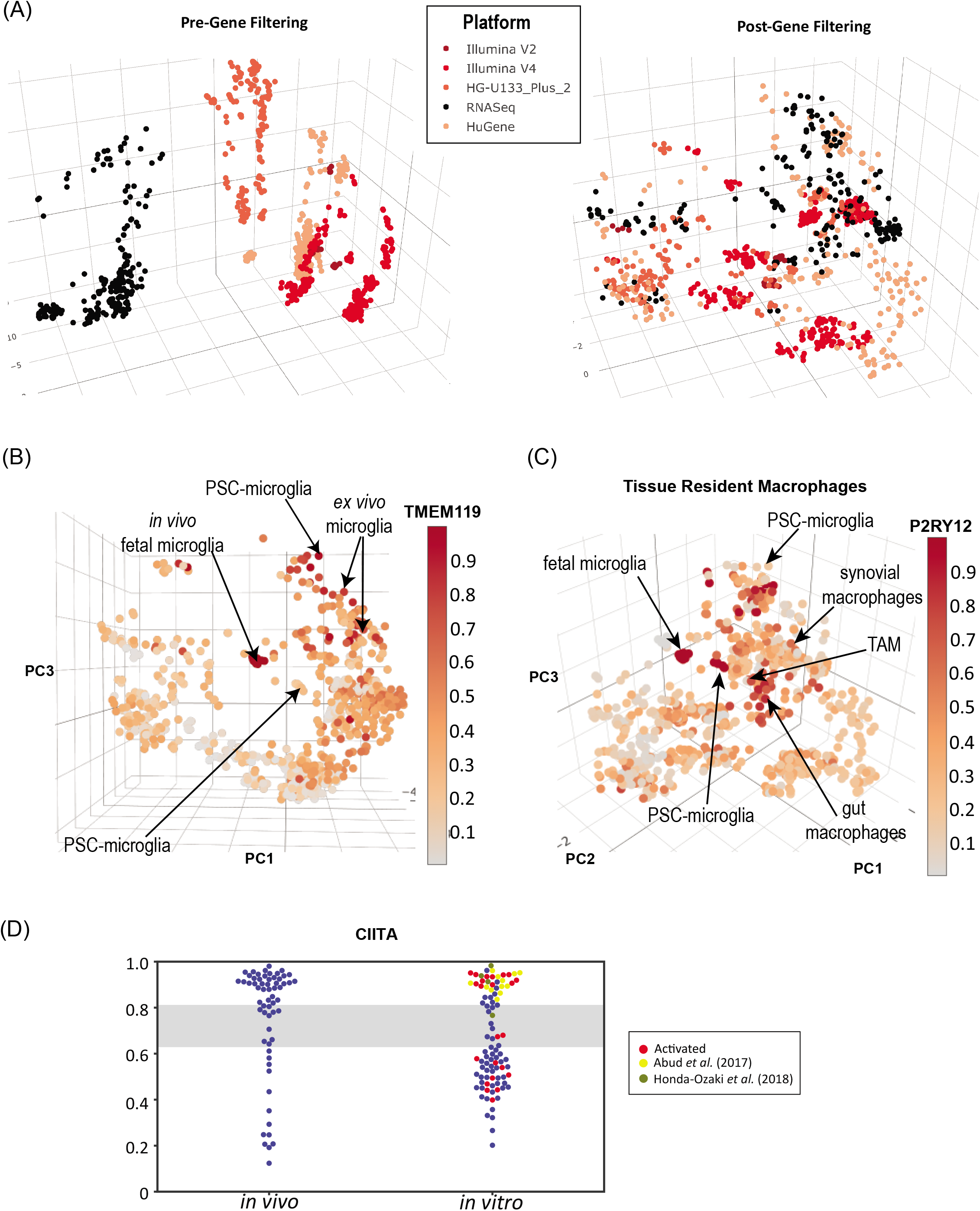
A reference atlas and resource for human myeloid biology. Related to Figure 1. (A) Pre-gene filtering (left) and Post-gene filtering (right) atlas coloured by platform: red various microarray platforms, black RNAseq platforms. (B) atlas coloured by ranked expression of TMEM119 (Scale bar: high ranked expression (dark red) to low ranked expression (grey) (C) atlas coloured by ranked expression of P2RY12 (Scale bar: high ranked expression (dark red) to low ranked expression (grey) (D) Ranked expression (Y-axis) of Class II transactivator (CIITA) *in vivo* versus *in vitro*-derived macrophages (gut, synovial, kupffer, microglia, macrophage). Red – activated, yellow – (Abud et al., 2017) microglia samples, khaki-Honda-Ozaki et al. (2018) macrophage samples.

**Figure S2:**
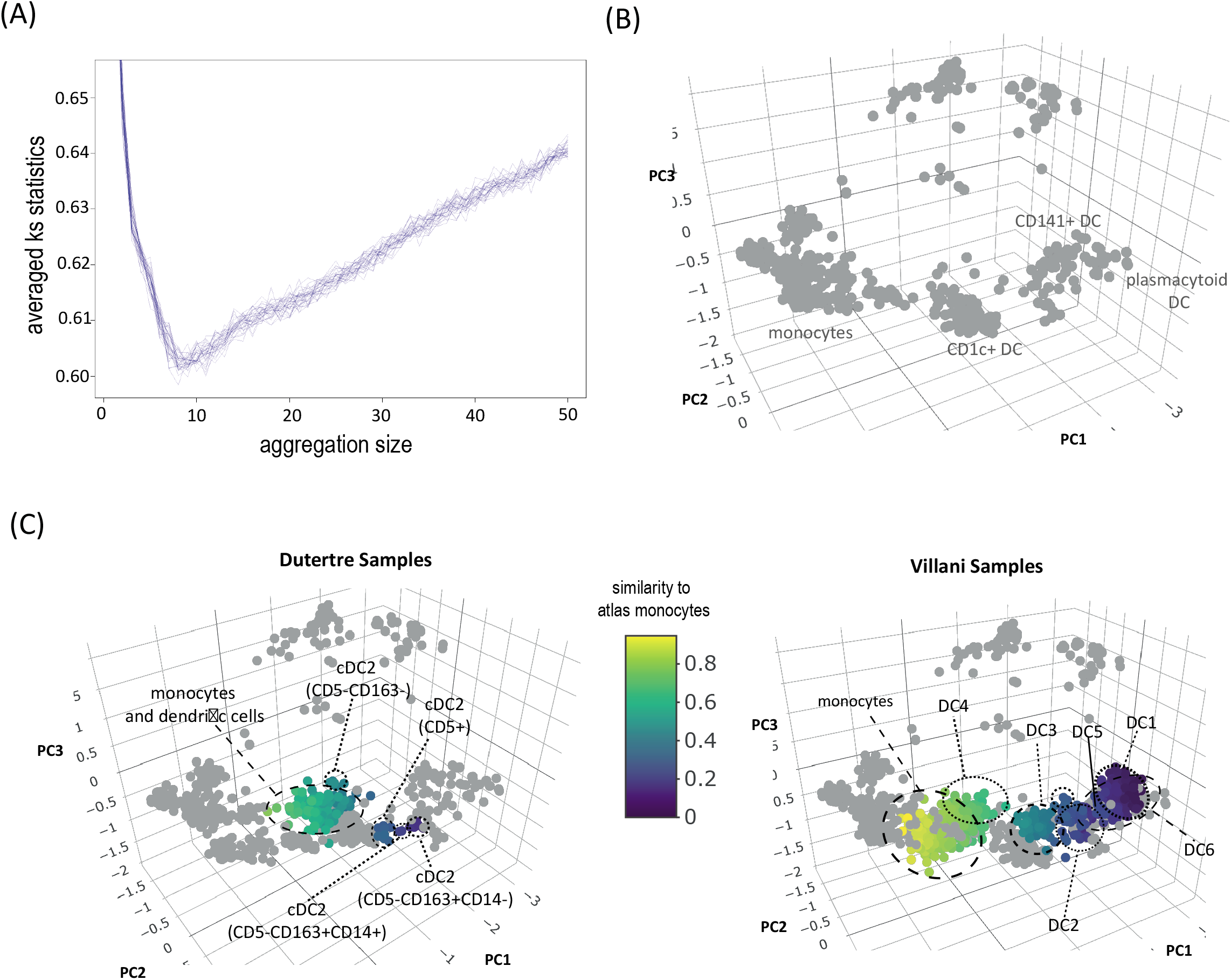
Single cell aggregation and projection. Related to Figure 2. (A) Kolmogorov–Smirnov (KS) statistics (y-axis) to assess the difference in gene expression distribution between pseudo-bulk single cells DC6 and bulk sample plasmacytoid dendritic cells from the atlas, with respect to the number of single cells that are aggregated (x-axis). Each line indicates one of thirty random sub-samplings with replacement trial. KS statistics are calculated on each gene and averaged across all genes. A minimum KS statistic is obtained when aggregating 8 cells. (B) atlas cell types before single cell projection (C) Single cell projection of (Dutertre et al., 2019) and (Villani et al., 2017) samples onto the atlas where a 8 cells are aggregated based on (A).

**Figure S3:**
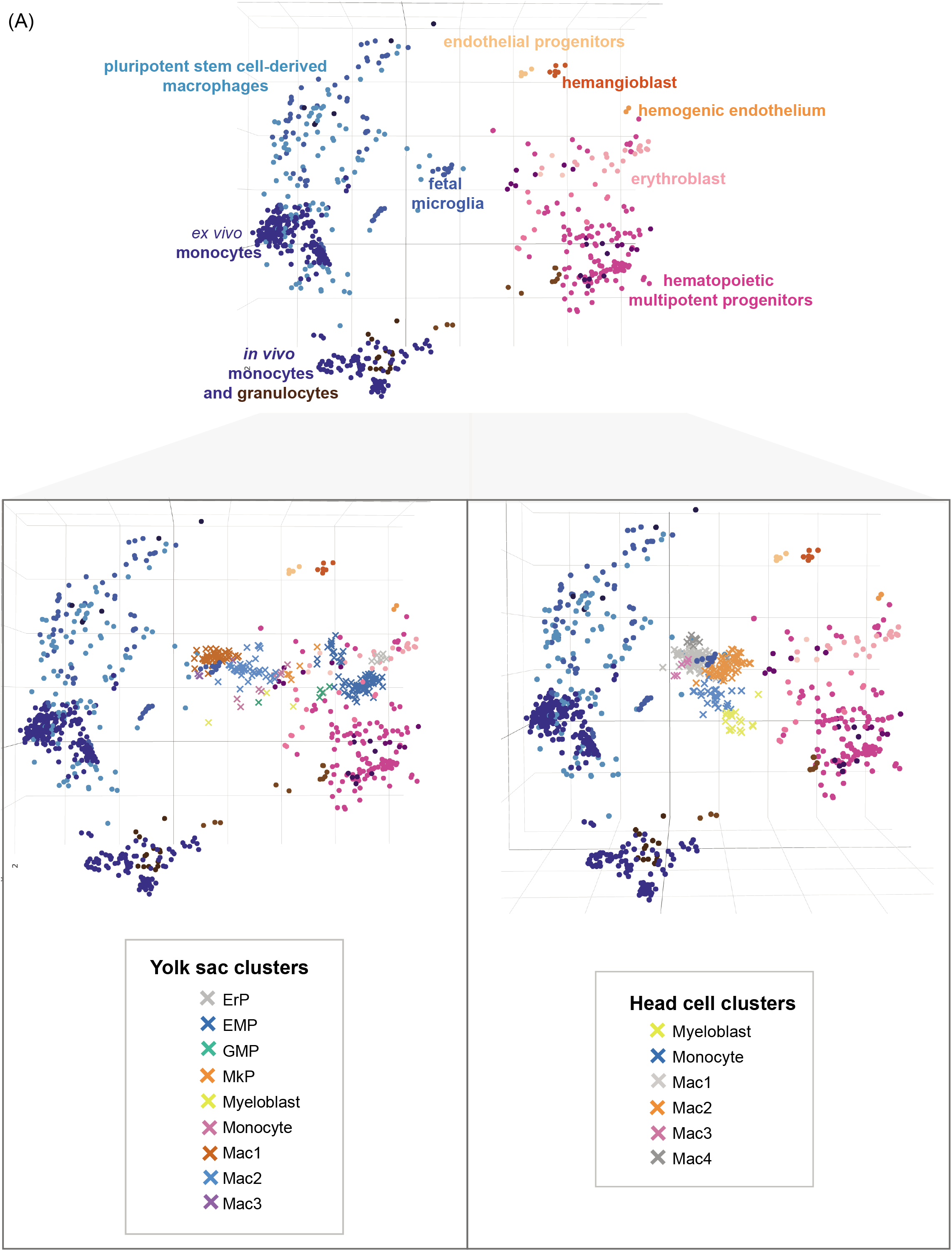
Fetal Ontogeny. Related to Figure 4. atlas coloured by cell type with (Bian et al., 2020) projection of single cell data from human fetal yolk sac and head.

## Supplemental Tables

**Table S1: Datasets and samples to compile atlas and single cell projection. Related to Figure 1.** Tissue resident macrophages and dendritic cells from peripheral blood, spleen, thymus, joint, lung, gut, brain and liver. Samples also included monocytes from peripheral and cord blood, as well as *in vitro* differentiated DCs from cord blood progenitors or monocyte-derived macrophages. Columns include dataset accession ID, platform, Stemformatics Dataset ID, number of samples, tier categorization, cell type and relevant tissue/organism part.

**Table S2: *in vivo* vs. *ex vivo* vs. *in vitro* dendritic cells. Related to Figure 2.** Comparison of gene expression of *in vivo* (n=145), *ex vivo* (n=17) and *in vitro*- (n=57) derived dendritic cells. Columns refer to gene symbols, P-values re-calculated by Mann-WhitneyWilcoxon rank-sum test, mean and standard deviation.

**Table S3: *in vivo* vs. *ex vivo* monocytes. Related to Figure 3.** Comparison of gene expression of *ex vivo* (n=171) and *in vivo* (n=107) monocytes. Columns refer to gene symbols, P-values re-calculated by Mann-Whitney-Wilcoxon rank-sum test, mean and standard deviation.

**Table S4: *in vivo* vs. *ex vivo* vs. *in vitro* macrophages. Related to Figure 4 and Figure 5.** Comparison of gene expression of *in vivo* (n=61), *ex vivo* (n=26), *in vitro*- (n=96) derived macrophages (gut, synovial, kupffer, microglia, macrophage). Columns refer to gene symbols, P-values recalculated by Mann-Whitney-Wilcoxon rank-sum test, mean and standard deviation.

**Table S5: *in vivo* vs. *ex vivo* vs. *in vitro* microglia. Related to Figure 4 and Figure 5.** Comparison of gene expression of in vivo (n=10), ex vivo (n=21) and in vitro- (n=43) derived microglia. Columns refer to gene symbols, P-values re-calculated by Mann-Whitney-Wilcoxon rank-sum test, mean and standard deviation.

**Table S6:** Gene-Set Enrichment Analysis. Related to Figure 6. Table of the top 10 Reactome pathways enriched in genes highly correlated with *in vitro*-derived macrophage spread. Enrichment: number of genes in the list/number of genes in that pathway (False Discovery Rate-value). Genes: multiple entries assigned to the same gene indicated by underlining of gene symbol with UniProt accessions in brackets.

| Table 1 |  |  |
| --- | --- | --- |
| Pathway | Enrichment | Genes |
| Collagen biosynthesis and modifying enzymes | 11/76 (3.76e-09) | ADAMTS3, COL1A2, COL4A2, SERPINH1, COL14A1, COL3A1, COL4A5, COL1A1, <u>COL4A1</u> (P02462, Q03692), COL5A1 |
| Extracellular matrix organization | 18/329 (4.48e-09) | ADAMTS1, COL1A1, <u>COL4A1</u> (P02462, Q03692), COL5A1, LTBP1, PTPRS, ADAMTS3, COL1A2, COL4A2, KDR, LUM, SERPINH1, COL14A1, COL3A1, COL4A5, LAMB1, MFAP4 |
| Collagen chain trimerization | 9/44 (4.81e-09) | COL14A1, COL3A1, COL4A5, COL1A1, <u>COL4A1</u> (P02462, Q03692), COL5A1, COL1A2, COL4A2 |
| Collagen formation | 11/104 (2.39e-08) | ADAMTS3, COL1A2, COL4A2, SERPINH1, COL14A1, COL3A1, COL4A5, COL1A1, <u>COL4A1</u> (P02462, Q03692), COL5A1 |
| ECM proteoglycans | 10/79 (2.39e-08) | COL1A1, COL4A1, COL5A1, PTPRS, COL1A2, COL4A2, LAMB1, COL3A1, COL4A5, LUM |
| Non-integrin membrane-ECM interactions | 9/61 (3.83e-08) | COL1A1, <u>COL4A1</u> (P02462, Q03692), COL5A1, COL1A2, COL4A2, LAMB1, COL3A1, COL4A5 |
| Integrin cell surface interactions | 10/86 (3.83e-08) | COL1A1, <u>COL4A1</u> (P02462, Q03692), COL5A1, COL1A2, COL4A2, KDR, COL3A1, COL4A5, LUM |
| Assembly of collagen fibrils and other multimeric structures | 9/67 (6.95e-08) | COL14A1, COL3A1, COL4A5, COL1A1, <u>COL4A1</u> (P02462, Q03692), COL5A1, COL1A2, COL4A2 |
| Collagen degradation | 9/69 (7.93e-08) | COL14A1, COL3A1, COL4A5, COL1A1, <u>COL4A1</u> (P02462, Q03692), COL5A1, COL1A2, COL4A2 |
| Degradation of the extracellular matrix | 11/148 (3.73e-07) | ADAMTS1, COL1A2, COL4A2, LAMB1, COL14A1, COL3A1, COL4A5, COL1A1, <u>COL4A1</u> (P02462, Q03692), COL5A1 |

## Supplemental Video

Active engulfment and clearance of cells by pluripotent stem cell-derived macrophages.

## Methods

Atlas formation was developed as described in (Angel et al., 2020) and is comprised of 44 datasets, 901 samples and 3757 genes. Mapping, and analysis of microarray and RNA sequencing datasets were conducted in the Stemformatics platform. Scripts are available for download from the Stemformatics BitBucket (Choi et al., 2019a)). All datasets and relevant samples (Supplementary Table 6) passed stringent quality control checks required for hosting on the stemformatics platform. Quality control checks include evaluation of library quality, and inclusion of replicates associated with experimental design. These datasets were either already hosted on stemformatics, or were downloaded from public depositories and processed through the stemformatics pipeline for inclusion.

### Platform Effect Analysis and Gene Selection for PCA

This method assesses each gene independently for a dependence upon experimental platform. The initial step is to transform expression values from RNA Sequencing and Microarray into percentile values. The second step uses a univariate estimate of gene platform dependence and then selects genes with a small ratio of platform dependent variance to total variance. These genes are used to run the PCA (3757 genes passed this cut). The threshold for gene selection is empirically determined by using the Kruskal Wallis H Test to assess the difference in platform expression distribution for each principal component. A platform variance fraction of 0.2 is found to remove the platform effect for the first three principal component. For further details, please refer to (Angel et al., 2020).

### Quantification and Statistical Analysis

P-values were re-calculated using the Mann-Whitney-Wilcoxon rank-sum test (two-sided). This was implemented via the python (version 3.7.5) SciPy package (version 1.3.1)(Virtanen et al., 2019)Multiple testing over the set of genes was accounted for with Bonferroni correction implemented in the statsmodels package (Seabold and Perktold, 2010)

### Pseudo-bulk samples from Villani’s Single Cell Data

Single cells were aggregated to form pseudo-bulk samples to mitigate library size differences between single cell and bulk data, and to project samples onto the atlas. Each group of (known) cell type in the single cell data was randomly sampled for *k* single cells with replacement. Aggregation consisted in summing up their expression profiles. k was determined by the number of sampling (i.e. how many pseud-bulk samples for each cell type) as half of the cell type’s population size. The optimum aggregation size *k* was investigated by evaluating the similarity in distribution between the atlas’ plasmacytoid dendritic cells and the aggregated pseudo-bulk DC6 samples using Kolmogorov–Smirnov (KS) statistic *D* averaged across all genes. KS statistic measured the difference between the empirical cumulative distribution functions of two groups of samples; the smaller the value, the closer the aggregated DC6 resembled the reference transcriptional profiles from the pDC in the bulk atlas. A minimum *D* value was obtained for *k* = 8 across 30 iterations (Supplementary Figure 1). Similar results were obtained for other cell types. Thus, every pseudo-bulk samples were aggregated from 8 single cells.

### Capybara Cell Score

Capybara (Kong et al., 2020)cell scores was used to measure cell identities continuum of the pseudo-bulk samples using the atlas as the reference. Capybara cell scores were calculated by performing restricted linear regression of reference samples on each of the pseudo-bulk samples’ expression profiles. Denote *y_i_* the expression profile of the *i^th^* pseudo-bulk sample of length *G*, where *G* represents the total number of genes in the data, and *X* a (*G*X*T*) the reference matrix, where *T* represents the number of known cell types of interest. We considered 5 cell types: Dendritic cell, Monocyte, CD141+ dendritic cell, CD1c+ dendritic cell and Plasmacytoid dendritic cells. *X* is obtained by averaging the expression profiles of the Stemformatics myeloid samples according to their cell types. Capybara solves the optimization problem

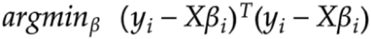

under the constraint that

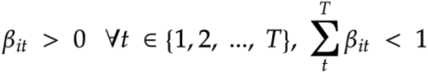

where *β_it_* is a regression coefficient, or cell score, for each pseudo-bulk sample *i* and each atlas cell type *t*. The cell score is obtained using quadratic programming implemented with R (version 3.6.2) package *quadprog (version 1.5-8)*(Turlach and Weingessel, 2019).

### Enrichment analysis and Protein-Protein Network

An enrichment analysis was conducted on the top 92 genes ranked by Pearson correlation (?0.7) along the upward axis including in vitro-derived cells. Enriched pathways were identified using these genes at Reactome (Fabregat et al., 2018) and significance ranked by p-value/false discovery rate. A protein-protein network was generated using the top 92 genes on STRING-DB (Szklarczyk et al., 2019). Disconnected nodes not shown.

### Pluripotent Stem Cell Differentiation

Stem cell work was carried out in accordance with The University of Melbourne ethics committee HREC (approval 1851831). Stem cell lines used were PB001.1 (Vlahos et al., 2019), a kind gift from the Stem Cell Core Facility at the Murdoch Children’s Research Institute, and HDF51((Jones et al., 2013); RRID:CVCL_UF42) was kindly provided to ALL by Prof. Jeanne Loring (The Scripps Research Institute, CA, USA). Human pluripotent stem cells were differentiated into macrophages based on protocol described by ((Joshi et al., 2019; Ng et al., 2008)), with modifications. Modifications were as follows: embryoid bodies were kept in rotational cultures without transference to matrigel plates for adherence, and the collection of progenitors from week 2 were immediately re-suspended in RPMI-1640 containing L-Glutamine (Life Technologies) and 10% Fetal Bovine Serum for macrophage differentiation (see macrophage differentiation).

### Monocyte isolation from peripheral blood

Buffy Coat was obtained from the Australian Red Cross Blood Service in accordance with The University of Melbourne ethics committee HREC (approval 1646608). The blood was diluted with PBS at a 1:3 dilution and underlayed with Ficoll-Hypaque. The underlayed blood samples were centrifuged at 350g for 30 minutes at 24°C with no brake. Peripheral blood mononuclear cells were isolated from the interphase and washed twice by using MACs buffer (DPBS, 0.5% heat inactivated Fetal Bovine Serum, 2mM EDTA) and centrifuging at 400g for 5 minutes at 4°C. Cells were centrifuged at 400g for 5 minutes at 4°C and resuspended in 40μl MACs buffer per 107 cells. Monocytes were positively selected by a magnetic field using Human CD14 MicroBeads (MACS Miltenyi Biotec) and LS Columns (Miltenybiotec). These cells were plated for macrophage differentiation (see Macrophage differentiation

### Macrophage differentiation

Monocytes/progenitor cells were cultured in tissue-culture treated 6 well plates. Cells were cultured in RPMI-1640 medium containing L-Glutamine (Life Technologies) with 10% Fetal Bovine Serum and 100ng/ml recombinant Human M-CSF (R&D Systems; 216-MC) for 5 days. Media changes were carried out on day 4.

### Flow Cytometry

HMDM and PSCM were collected and centrifuged at 400g for 5 minutes. Supernatant was aspirated and 5μl mouse serum was added to ‘dry’ pellets for 5 minutes on ice. Cells were resuspended in FACS Buffer (Hanks Balanced Salt Solution, 0.5% Human Serum Albumin) and stained with CD14 or matched isotype control antibodies on ice for 20 minutes then washed twice (3ml FACS Buffer, spun at 400g, 5 minutes). Resuspended cells were fixed with 4% paraformaldehyde (PFA) for 15 minutes at room temperature. PFA was washed out and cells rinsed twice in FACS buffer before resuspending in PBS and stored overnight at 4°C. Fixed cells were permeabilized with 0.1% Triton X-100 (in 1XPBS) for 10 minutes at room temperature and washed twice. Blocking buffer (0.3M glycine buffer, 10% Goat Serum, 1XPBS) was added to ‘dry’ pellets on ice for 1 hour. Cells were stained with antibodies to Type I Collagen or matched isotype control on ice for 20 minutes. Cells were washed twice then incubated with secondary antibody for 20 minutes on ice in the dark. Cells were then washed twice and resuspended for analysis on a CytoFLEX S flow cytometer (Beckman Coulter, Brea, CA) using CytExpert acquisition software. Post-acquisition analysis was performed with FCS Express 7 flow cytometry software.

| Flow Cytometry antibodies |  |
| --- | --- |
| CD14 | Brilliant Violet 421™ anti-Human CD14 (BioLegend: Cat. No. 325628; RRID:AB_2563296 ) |
| CD14 Isotype | Brilliant Violet 421™ Mouse IgG1 (BioLegend: Cat. No. 400157; RRID:AB_10897939) |
| Type I Collagen Primary antibody | Rabbit Anti-Collagen I antibody (Abcam: Cat. No. ab264074) |
| Secondary Antibody | Goat Anti-Rabbit IgG H&L (Alexa Fluor®488) (Abcam: Cat. No. ab150077; RRID:AB_2630356) |
| Type I Collagen Isotype | Rabbit IgG monoclonal Isotype Control (Abcam: Cat. No. Ab172730; RRID:AB_2687931) |

### Stimulation Assay

On day 5 of differentiation, one well containing peripheral blood monocyte- or human pluripotent stem cell-derived macrophages were stimulated with 10ng/ml Lipopolysaccharide (LPS) (Sigma-Aldrich; *Salmonella enterica* serotype minnesota) for 2 hours. After stimulation period, media was aspirated, and the wells were washed twice with PBS (Ca2+Mg2+ free). before cell lysis using 2-mercaptoethanol (Sigma-Aldrich) and RNeasy Plus Lysis Buffer (Qiagen). Samples were placed into Eppendorf’s and stored at −80°C before RNA extraction.

### RNA extraction

Total RNA was isolated using the RNeasy^®^ Plus Mini Kit (Qiagen) according to manufacturer’s instructions. In summary: for the removal of genomic DNA, samples were placed into gDNA columns and centrifuged for 30 seconds at 8000g. Ethanol (70%) was mixed with the flow through and samples were transferred to RNeasy spin columns. The columns were centrifuged for 15 seconds at 8000g. Buffer RW1 was then added to the columns and columns were centrifuged for 15 seconds at 8000g. Buffer RPE was added to the columns and columns were centrifuged for 15 seconds. Buffer RPE was again added to the columns with centrifugation at 8000g for 2 minutes. Columns were then placed into new collection tubes and centrifuged at full speed for 1 minute to dry the membrane. RNase-free water was then added directly onto the column membrane and columns placed into Eppendorf’s and centrifuged at 8000g for 1 minute to collect RNA. RNA quality and quantity were determined using a Tapestation (Agilent Technologies 2200). Samples were stored at −80°C. Zymo Research RNA Clean & Concentrator-25 Kit was used to pool replicates (from the same donor) together and elute in elute into smaller volume with maximum concentration.

### RNA sequencing

RNA samples were processed by the Ramaciotti Centre for Genomics (University of New South Wales; Sydney). Illumina Novaseq_6000 was used for mRNA-sequencing.

### Data and Code Availability

mRNA-sequencing data is available through accession GSE150893. All public accessions are listed in Supplementary Table 6. Stemformatics code is publicly available at bitbucket.org/stemformatics. Atlas code is available at bitbucket.org/stemformatics/s4m_pyramid/src/master/scripts/atlas.py.

### Graphing software and Illustration

Graphs for mRNA-seq gene expression were generated using Graphpad Prism. Violin plots were generated through the www.stemformatics.org platform. Schematic Figure illustrations created with BioRender.com

